# Caffeine and MDMA (ecstasy) exacerbate ER stress triggered by hyperthermia

**DOI:** 10.1101/2022.01.14.476356

**Authors:** Kathleen A. Trychta, Brandon K. Harvey

## Abstract

Drugs of abuse can cause local and systemic hyperthermia, a known trigger of endoplasmic reticulum (ER) stress and the unfolded protein response (UPR). Another trigger of ER stress and UPR is ER calcium depletion which causes ER exodosis, the secretion of ER resident proteins. Club drugs such as 3,4-methylenedioxymethamphetamine (MDMA, ‘ecstasy’) can create hyperthermic conditions in the brain and cause toxicity that is affected by the environmental temperature and the presence of other drugs, such as caffeine. Here we examine the secretion of ER resident proteins and activation of the UPR under combined exposure to MDMA and caffeine in a cellular model of hyperthermia. We show that hyperthermia triggers the secretion of normally ER resident proteins and that this aberrant protein secretion is potentiated by the presence of MDMA, caffeine, or a combination of the two drugs. Hyperthermia activates the UPR but the addition of MDMA or caffeine does not alter canonical UPR gene expression despite the drug effects on ER exodosis of UPR-related proteins. One exception was increased BiP/Grp78 mRNA levels in MDMA-treated cells exposed to hyperthermia. These findings suggest that club drug use under hyperthermic conditions exacerbates disruption of ER proteostasis contributing to cellular toxicity.

**Highlights:**

1. ER resident proteins are redistributed into the extracellular space in response to hyperthermia and caffeine and MDMA further enhance this secretion.
2. Stabilizing ER calcium and overexpressing KDEL receptors reduces ER resident protein secretion following hyperthermia.
3. Hyperthermia triggers a UPR response with MDMA augmenting BiP expression in hyperthermic conditions.

## Introduction

Club drugs or “rave” drugs are a class of recreational drugs associated with the location of usage, such as nightclubs, raves, and dance parties. In these environments, the combination of a warm atmosphere, overcrowding, and dancing contribute to elevated body temperature or hyperthermia, which can negatively impact the body’s reaction to substances of abuse. For example, the combination of hyperthermic environmental conditions and one of the most widely used club drugs, 3,4-methylenedioxymethamphetamine (MDMA, ‘ecstasy’), can be lethal in humans^1, 2^. Moreover, MDMA is associated with drug-induced hyperthermia and consumption of the drug can itself cause hyperthermia in humans ^3-6^. The potentiation of MDMA-induced hyperthermia by social interaction and warm ambient temperature has been recreated in rodent models^7^. Consumption of other stimulants, such as caffeine, has been shown to promote hyperthermia and toxicity associated with MDMA^8^. Caffeine is commonly present in drinks consumed with club drugs and is also used as a drug additive. The interplay between increased ambient temperature, drug usage, and cellular proteostasis has not been well documented.

Human body temperature is normally maintained at 37°C with fluctuations of only a degree or two being well tolerated and temperatures that reach 40°C or higher generally constituting hyperthermia. The central nervous system is vulnerable to hyperthermia and increased temperatures have been linked to cognitive dysfunction, seizures, and loss of consciousness as well as exacerbating poor outcomes following brain injury (i.e. traumatic brain injury, subarachnoid hemorrhage, stroke, substance abuse)^3, 9-11^. At a cellular level, as temperatures rise above 37°C the amount of unfolded proteins, protein aggregates, and denatured proteins increases^12, 13^. Under normal conditions when the cell detects misfolded proteins the unfolded protein response (UPR) is activated to promote the degradation of misfolded proteins, diminish overall protein production, and upregulate molecular chaperones to restore protein folding ^14, 15^. While hyperthermia triggers the UPR, this adaptive response does not always prevent cell death ^16-19^. Furthermore, elevated temperatures lead to decreased chaperone activity and some chaperones can function as proteases at increased temperatures, preventing the ER from re-establishing protein homeostasis^20^. Hyperthermic disruptions to ER homeostasis have the potential to affect intrinsic ER functions like protein processing and trafficking, lipid and carbohydrate metabolism, and drug detoxification as these ER processes rely on the presence and activity of ER luminal proteins ^21-24^.

Studies of ER resident proteins have identified carboxy-terminal peptide sequences referred to as ER retention sequences (ERS) that localize proteins to the ER lumen^25-29^. If an ERS-containing protein escapes the ER lumen the ERS sequence is recognized by KDEL receptors in the Golgi and the KDEL receptors interact with the protein to facilitate its retrograde transport back to the ER ^25, 30^. Under certain stress conditions, ERS-containing proteins are secreted from the cell^31-35^. A redistribution of ERS proteins en masse from the ER lumen to the extracellular space was observed in response to both ER calcium depletion and oxygen-glucose deprivation^36^. This phenomenon, termed ER exodosis, refers to the departure of a resident protein from its organelle under pathophysiological conditions. Given previously described links between hyperthermia and both ER stress and changes in cellular calcium, hyperthermia represents a putative trigger of ER exodosis^16, 37^.

Here we investigate the effects of environment-induced hyperthermia on ER proteostasis in the presence and absence of MDMA and caffeine. We show for the first time that hyperthermia causes the secretion of ER resident proteins, or ER exodosis. We find that both MDMA and caffeine exacerbate ER exodosis under hyperthermic conditions in both a human neuronal cell line and rodent primary cortical neurons. Our findings identify a cellular mechanism impacted by hyperthermia and exposure to the drugs MDMA and caffeine. The therapeutic potential of preventing ER exodosis in the treatment of club drug toxicity and hyperthermia is discussed.

## Results

### Hyperthermia triggers the secretion of ER resident proteins

In order to study the secretion of ER resident proteins in a model of hyperthermia we used a *Gaussia* luciferase (GLuc) protein with an ERS. This GLuc with an ASARTDL (Ala-Ser-Ala-Arg-Thr-Asp-Leu) C-terminal ERS was previously shown to localize to the ER lumen and act as a marker of ER exodosis^35, 36^. Using a human SH-SY5Y neuroblastoma cell line that stably expresses the GLuc-ASARTDL reporter as well as a cell line expressing a constitutively secreted GLuc reporter with no ERS (GLuc-Untagged) we discovered that following hyperthermia there was increased secretion of GLuc-ASARTDL whereas there was no change in GLuc-Untagged secretion (Fig. 1A-B). Prolonged exposure to hyperthermia caused a decrease in cell viability (indicated by ATP levels) in both cell lines consistent with results seen previously in SH-SY5Y (Supp. Fig. 1A-B)^35, 38^. To examine whether the increased secretion of GLuc-ASARTDL was reflective of the behavior of endogenous ER resident proteins, we examined the extracellular levels of three different endogenous ERS-containing proteins (MANF, PDI, esterases). For each of these ER resident proteins, extracellular protein levels were increased following hyperthermia suggesting that hyperthermia leads to ER exodosis in which there is a mass redistribution of ERS-containing ER resident proteins into the extracellular space following an insult (Fig. 1C-E). We further examined the effects of hyperthermia in rat primary cortical neurons (PCNs) by transducing PCNs with an AAV vector expressing GLuc-ASARTDL. Similar to what was observed in SH-SY5Y cells, an increase in extracellular GLuc-ASARTDL was observed following hyperthermia in PCN (Supp. Fig. 1C). No such increase in GLuc was observed in PCNs transduced with GLuc-Untagged although cells exhibited comparable hyperthermia-induced decreases in ATP levels regardless of the reporter used (Supp. Fig. 1D-F).

**Figure 1:**
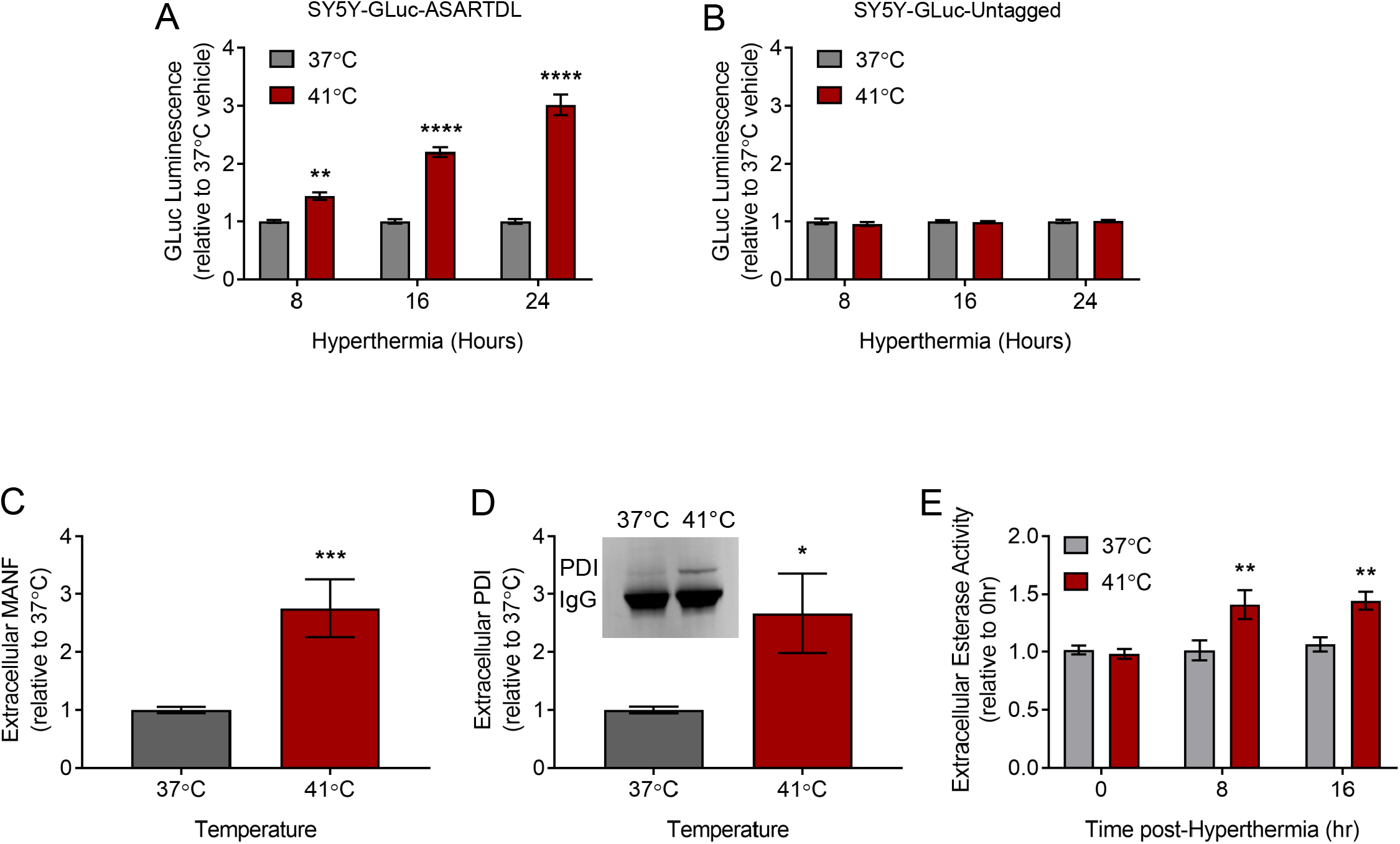
ER resident protein secretion is triggered by hyperthermia. (A-B) GLuc activity in the media from SH-SY5Y cells stably expressing (A) GLuc-ASARTDL or (B) GLuc-Untagged after an 8 h, 16 h, or 24 h incubation at 37°C or 41°C (mean ± SEM, n=16, two-way ANOVA with Sidak’s multiple comparisons, **p<0.01 and ****p<0.0001 37°C vs. 41°C). (C-D) SH-SY5Y cells were exposed to 37°C or 41°C for 24 h. (C) Fold change in extracellular MANF from SH-SY5Y cells following a 24 h incubation at 37°C or 41°C (mean ± SEM, n=5, t-test, ***p<0.001). (D) Fold change in immunoprecipitated PDI (representative blot shown) from media of SH-SY5Y cells following incubation at 37°C or 41°C (mean ± SEM, n=3, t-test, *p<0.05). (E) SH-SY5Y cells were exposed to 37°C or 41°C for 8 h or 16 h. Fluorescent esterase assay of cell culture media presented as a fold change in extracellular esterase activity (mean ± SEM, n=12, two-way ANOVA with Sidak’s multiple comparisons, **p<0.01 37°C vs. 41°C).

### Manipulating ER calcium alters hyperthermia-induced ER exodosis

Hyperthermia has a complex etiology with some links to changes in intracellular calcium^39, 40^. In order for the ER to carry out its myriad of functions, luminal calcium concentrations must be tightly controlled. Under basal conditions, the sarco-/endoplasmic reticulum calcium ATPase (SERCA) pumps calcium from the cytoplasm into the ER while the ryanodine receptor (RyR) and inositol 1,4,5-triphosphate receptor (IP3R) allow for calcium efflux ^41-43^. Reductions in ER calcium are associated with ER resident protein secretion^32-34, 36^. In our hyperthermia model, treatment with thapsigargin, a SERCA inhibitor, potentiated the release of GLuc-ASARTDL in hyperthermic conditions (Supp. Fig. 2A). To determine whether this effect occurred following other forms of ER stress we treated cells with tunicamycin. Both thapsigargin and tunicamycin are known to cause ER stress with thapsigargin inhibiting ER calcium uptake and tunicamycin acting as an N-linked glycosylation inhibitor and were used at doses previously shown to induce comparable levels of ER stress^34, 44, 45^. Only thapsigargin enhanced GLuc-ASARTDL secretion in hyperthermic conditions (Supp. Fig. 2B). Treatment with tunicamycin caused no change in GLuc-ASARTDL levels when compared to vehicle treated cells suggesting that in hyperthermic conditions GLuc-ASARTDL secretion is further increased following changes in ER calcium but not changes to ER stress in general (Fig. 2B).

**Figure 2:**
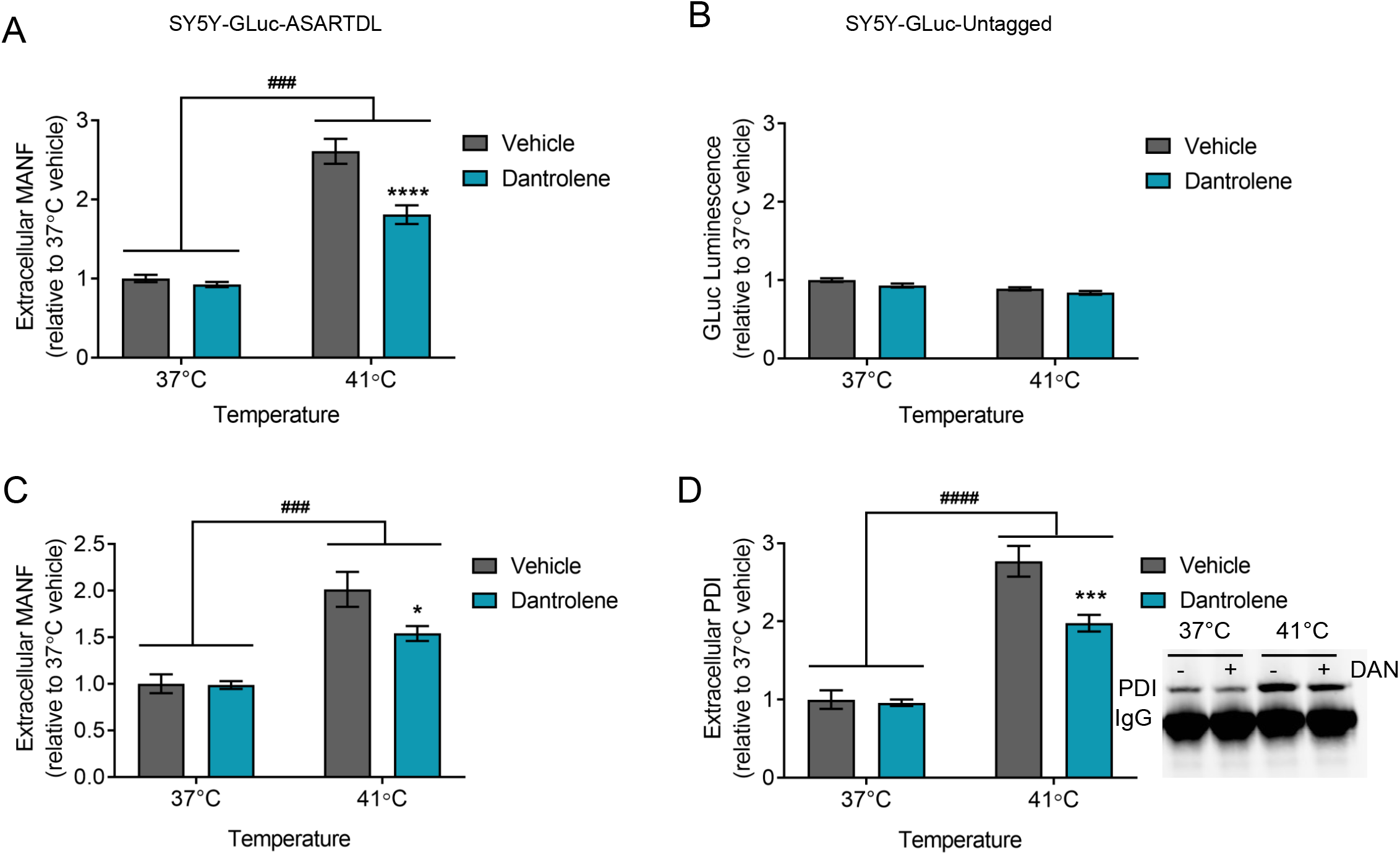
Blocking ER calcium efflux attenuates hyperthermia-induced ER resident protein secretion. (A-B) GLuc activity in the media from SH-SY5Y cells stably expressing either (A) GLuc-ASARTDL or (B) GLuc-Untagged after a 16 h pre-treatment with 3 μM dantrolene followed by a 24 h incubation at 37°C or 41°C (mean ± SEM, n=12, two-way ANOVA with Dunnett’s multiple comparisons, ^###^p<0.001 37°C vs. 41°C, ****p<0.0001 vehicle vs. dantrolene). (C) Fold change in extracellular MANF from SH-SY5Y cells following a 16 h pre-treatment with 3 μM dantrolene and 24 h exposure to 37°C or 41°C (mean ± SEM, n=3, two-way ANOVA with Sidak’s multiple comparisons, ^###^p<0.001 37°C vs. 41°C, *p<0.05 vehicle vs. dantrolene). (D) Fold change in immunoprecipitated PDI (representative blot shown) from the media of SH-SY5Y cells following a 16 h pre-treatment with 3 μM dantrolene and 24 h exposure to 37°C or 41°C (mean ± SEM, n=6, two-way ANOVA with Sidak’s multiple comparisons, ^####^p<0.0001 37°C vs. 41°C, ***p<0.001 vehicle vs. dantrolene).

In addition to altering calcium influx into the ER, we also examined how blocking calcium efflux from the ER affects ER exodosis. Dantrolene, a RyR antagonist, can attenuate ER calcium depletion and decrease ER resident protein secretion^34-36, 46^. In our model of hyperthermia, treating with dantrolene attenuated the secretion of GLuc-ASARTDL observed following hyperthermia, while having no effect on the secretion of GLuc-Untagged (Fig. 2A-B, Supp. Fig. 2C). Treatment with dantrolene did not affect cellular ATP levels (Supp. Fig. 2D). An examination of MANF and PDI, two endogenous ERS-containing proteins, found an increase in extracellular protein levels following hyperthermia which was diminished by treatment with dantrolene (Fig. 2C-D). The behavior of endogenous MANF and PDI mirrored the results seen with the GLuc-ASARTDL reporter, supporting a model in which hyperthermia leads to ER exodosis that can be partially blunted by stabilizing ER calcium. Despite the ability of dantrolene to attenuate hyperthermia-induced secretion in SH-SY5Y, dantrolene was unable to reduce GLuc-ASARTDL secretion in PCNs (Supp. Fig. 2E). As dantrolene primarily acts on the RyR1 and RyR3 isoforms of the RyR and neurons primarily express RyR2 we expanded our analysis to look at modulators of IP3R activity^47, 48^. Stabilizing ER calcium by inhibiting IP3R with 2-APB reduced extracellular GLuc levels in cells transduced with GLuc-ASARTDL and exposed to hyperthermia, while no effect was seen on cells transduced with GLuc-Untagged (Supp. Fig. 2F) ^49^. 2-APB did not significantly affect cell viability (Supp. Fig. 2G). Like the results seen with GLuc-ASARTDL, extracellular levels of the endogenous ERS-containing protein PDI were increased following hyperthermia with reduced levels observed in 2-APB treated cells (Supp. Fig. 2H).

### Hyperthermia affects the unfolded protein response

The unfolded protein response is an adaptive cellular response to ER stress that seeks to restore proteostasis and maintain cell viability. Previous studies indicate that hyperthermia provokes the UPR and we show here that incubation at 41°C results in upregulation of the UPR target genes BiP, ERdj4, and ASNS (Fig. 3A)^16, 18^. There appears to be a link between UPR activation and length of exposure to hyperthermic conditions as all three UPR target genes are upregulated at 8 h and 24 h, but only ERdj4 is upregulated at 4 h (Fig. 3A). ERdj4 is a target gene of the IRE1α/XBP1 prong of the UPR. As previous work from our lab demonstrated that ER exodosis is modulated by the KDEL receptor retrieval pathway and that KDELR2 and KDELR3 are UPR responsive genes regulated by XBP1 we sought to further examine these findings in our model of hyperthermia ^36^. In our model, inhibiting IRE1α kinase activity with KIRA6 potentiated hyperthermia-induced GLuc-ASARTDL secretion, but had no effect on GLuc-ASARTDL in normothermic conditions in both SH-SY5Y cells and PCNs (Fig. 3B, Supp. Fig. 3A)^50^. Treatment with KIRA6 did not increase extracellular GLuc levels in GLuc-Untagged cells (Supp. Fig. 2B-C). Despite the observed upregulation of ERdj4 and increased ER exodosis following treatment with KIRA6, mRNA levels of the KDEL receptors remained relatively unchanged with only KDELR1 being minimally upregulated following hyperthermia (Fig. 3C). However, knocking down KDEL receptors increased GLuc-ASARTDL and MANF release indicating that the KDEL receptors still play an important role in hyperthermia-induced ER exodosis (Fig. 3D, Supp. Fig. 3D). GLuc-Untagged secretion was not affected by KDEL receptor knockdown (Supp. Fig. 3E). Conversely, overexpressing KDEL receptors decreased GLuc-ASARTDL secretion following hyperthermia (Fig. 3E). Taken together these results indicate that hyperthermia triggers the UPR and that ER exodosis is sensitive to manipulations of the KDEL retrieval pathway.

**Figure 3:**
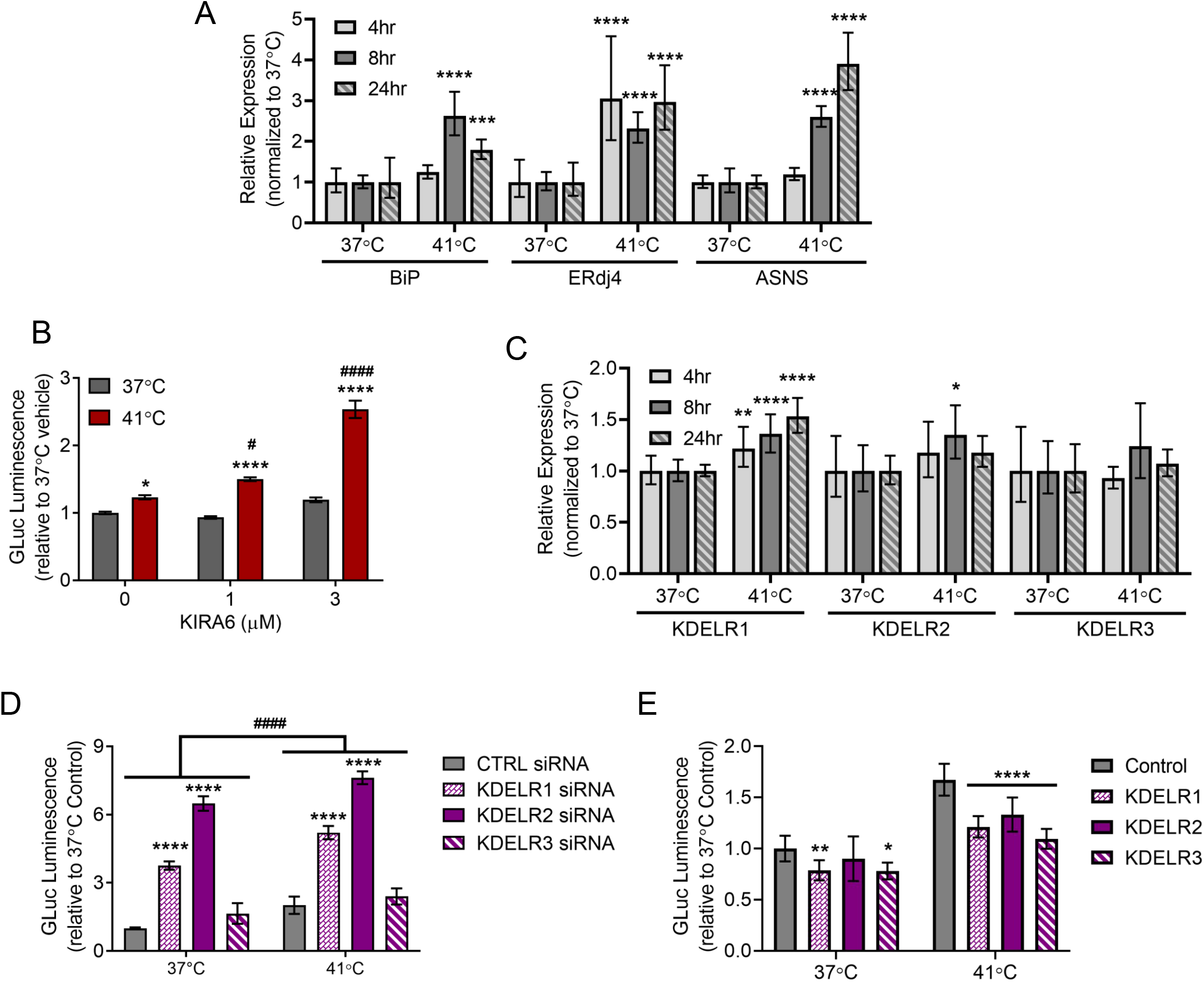
The UPR and KDEL receptors in hyperthermic conditions. (A) BiP, ERdj4, and ASNS mRNA levels analyzed with real-time RT-qPCR after a 4, 8, or 24 h exposure to 37°C or 41°C (2^−ddCq^ ± upper and lower limits, n=8, two-way ANOVA with Sidak’s multiple comparisons, ***p<0.001 and ****p<0.0001 37°C vs. 41°C). (B) GLuc activity in the media from SH-SY5Y cells stably expressing GLuc-ASARTDL after a 1 h treatment with vehicle or KIRA6 (1 μM or 3 μM) followed by a 24 h incubation at 37°C or 41°C (mean ± SEM, n=48, two-way ANOVA with Tukey’s multiple comparisons, *p<0.05 and ****p<0.0001 37°C vs. 41°C, ^##^p<0.01 and ^####^p<0.0001 vehicle vs. KIRA6). (C) KDELR1, KDELR2, and KDELR3 mRNA levels analyzed with real-time RT-qPCR after a 4, 8, or 24 h exposure to 37°C or 41°C (2^−ddCq^ ± upper and lower limits, n=8, two-way ANOVA with Sidak’s multiple comparisons, *p<0.05, **p<0.01, and ****p<0.0001 37°C vs. 41°C). (D) GLuc activity in media following transfection of SH-SY5Y cells stably expressing GLuc-ASARTDL with 10 nM KDEL receptor siRNA (mean ± SEM, n=12, two-way ANOVA with Dunnett’s multiple comparisons, ^####^p<0.0001 37°C vs. 41°C, ****p<0.0001 control vs. KDELR siRNA). (E) GLuc activity in media following transduction of SH-SY5Y cells stably expressing GLuc-ASARTDL with lentiviral KDEL receptors and a 24 h exposure to 37°C or 41°C (mean ± SEM, n=12, two-way ANOVA with Dunnett’s multiple comparisons, p<0.0001 37°C vs. 41°C, *p<0.05, **p<0.01, and ****p<0.0001 control vs. LV-KDELR).

### Caffeine and MDMA affect ER responses to hyperthermia

Caffeine is one of the most widely used drugs in the world and it is known to affect ER calcium by acting as a RyR agonist. We previously showed that GLuc-ASARTDL in the media was increased following a 72 h incubation with 5 and 10 mM caffeine ^35^. Therefore, we sought to examine the combined effects of a more acute caffeine treatment and hyperthermia on UPR activation and ER exodosis. As seen previously, hyperthermia triggered a UPR response in cells. Caffeine treatment did not augment UPR activation in hyperthermic cells, but did result in decreased BiP mRNA levels as well as increased ASNS mRNA levels at normal temperatures (Fig. 4A). Caffeine alone did significantly change GLuc-ASARTDL secretion, but caffeine did trigger a robust increase in secretion under hyperthermic conditions (Fig. 4B). GLuc-Untagged secretion was unaffected by hyperthermia or caffeine (Fig. 4B). Caffeine at the highest dose (10 mM) was associated with reduced cellular ATP levels (Supp. Fig. 4A). In accordance with the GLuc-ASARTDL data, both extracellular PDI and MANF were increased following hyperthermia and caffeine caused a significant, dose-dependent increase in extracellular levels of both endogenous ERS proteins (Fig. 4C-D). Similar findings were seen in PCNs with caffeine augmenting hyperthermia-induced ER exodosis and decreasing ATP levels (Supp. Fig. 4B-D). As demonstrated previously, treating with dantrolene reduced GLuc-ASARTDL secretion caused by hyperthermia, but only a minimal effect was seen when cells were exposed to both hyperthermia and caffeine perhaps because both caffeine and dantrolene act on the RyR or the effects of caffeine on other signaling pathways exacerbate hyperthermia-induced exodosis (Supp. Fig. 4E). GLuc-Untagged secretion was not affected by hyperthermia, caffeine, or dantrolene (Supp. Fig. 4F). Neither hyperthermia nor caffeine changed KDEL receptor mRNA levels, but overexpression of KDEL receptors inhibited hyperthermia and caffeine induced GLuc-ASARTDL secretion (Fig. 4E-F).

**Figure 4:**
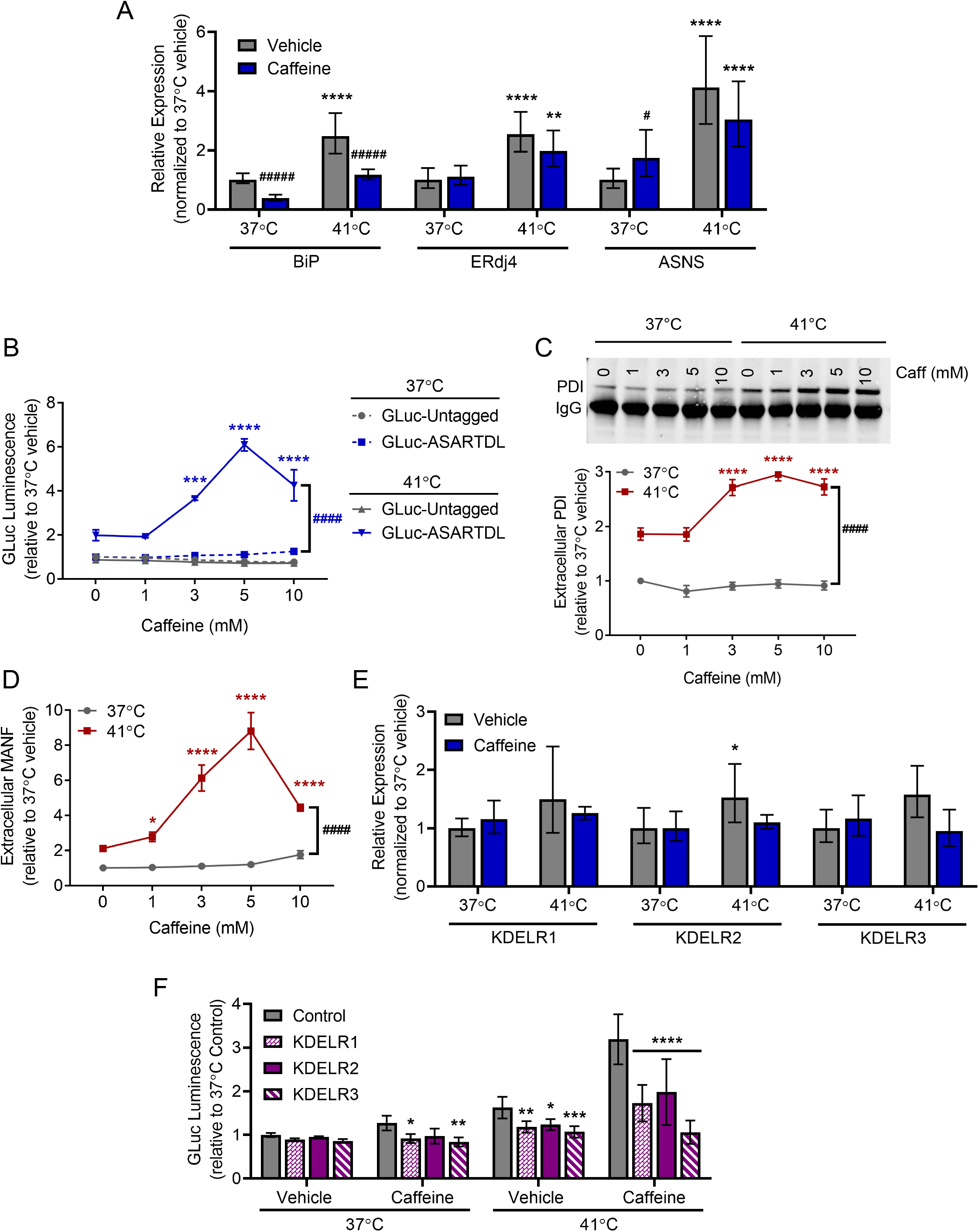
Caffeine changes cellular responses to hyperthermia. (A) BiP, ERdj4, and ASNS mRNA levels analyzed with real-time RT-qPCR after treatment with vehicle or 5 mM caffeine and an 8 h exposure to 37°C or 41°C (2^−ddCq^ ± upper and lower limits, n=8, two-way ANOVA with Sidak’s multiple comparisons, **p<0.01 and ****p<0.0001 37°C vs. 41°C, ^#^p<0.05, ^####^p<0.001 vehicle vs. caffeine). (B) GLuc activity in the media from SH-SY5Y cells stably expressing GLuc-ASARTDL or GLuc-Untagged after treatment with vehicle or caffeine and a 24 h incubation at 37°C or 41°C (mean ± SEM, n=26, two-way ANOVA with Tukey’s multiple comparisons, ^####^p<0.0001 37°C vs. 41°C, ***p<0.001 and ****p<0.0001 vehicle vs. caffeine). (C-D) Fold change in (C) immunoprecipitated PDI (representative blot shown) or (D) MANF in media from SH-SY5Y cells treated with vehicle or caffeine and incubated for 24 h at 37°C or 41°C (mean ± SEM, n=6, two-way ANOVA with Dunnett’s multiple comparisons, ^####^p<0.0001 37°C vs. 41°C, *p<0.05 and ****p<0.0001 vehicle vs. caffeine). (E) KDELR1, KDELR2, and KDELR3 mRNA levels analyzed with real-time RT-qPCR after treatment with vehicle or 5 mM caffeine and an 8 h exposure to 37°C or 41°C (2^−ddCq^ ± upper and lower limits, n=8, two-way ANOVA with Sidak’s multiple comparisons, *p<0.05 37°C vs. 41°C). (F) GLuc activity in media following transduction of SH-SY5Y cells stably expressing GLuc-ASARTDL with lentiviral KDEL receptors and treatment with vehicle or 5 mM caffeine and a 24 h exposure to 41°C (mean ± SEM, n=9, two-way ANOVA with Dunnett’s multiple comparisons, p<0.0001 37°C vs. 41°C, *p<0.05, **p<0.01, ***p<0.001, and ****p<0.0001 control vs. LV-KDELR).

3,4-methylenedioxymethamphetamine (MDMA) is a drug of abuse that can have more severe effects in certain environmental conditions, including elevated temperatures^51^. SH-SY5Y cells have been used previously to study the effects of MDMA, so we wanted to examine whether MDMA affected ER stress and exodosis^52, 53^. We observed a marked increase in BiP expression in hyperthermic conditions when MDMA was present, although ERdj4 and ASNS levels were not changed by the presence of MDMA (Fig. 5A). Following exposure to MDMA in hyperthermic conditions we found increased GLuc-ASARTDL secretion beyond that seen in vehicle conditions (Fig. 5B). MDMA did not change GLuc-ASARTDL secretion in normothermic controls and the control reporter, GLuc-Untagged, was not increased by hyperthermia or MDMA (Fig. 5B). MDMA also caused a decrease in cellular ATP at higher doses under hyperthermic conditions (Supp. Fig. 5A). Extracellular levels of the ERS-containing proteins PDI and MANF were increased following hyperthermia and were more highly secreted with increasing concentrations of MDMA suggesting that MDMA increases the secretion of ERS-containing proteins, but only in hyperthermic conditions (Fig. 5C-D). Studies in PCNs revealed a similar pattern with MDMA exacerbating ER resident protein secretion and decreasing cell viability at elevated temperatures (Supp. Fig. 5B-D). Stabilizing ER calcium by treating with dantrolene reduced GLuc-ASARTDL secretion resulting from hyperthermia and MDMA, while GLuc-Untagged secretion was unchanged (Supp. Fig. 5E-F). KDEL receptor expression was not changed by increased temperature or MDMA, but overexpression of KDEL receptors attenuated GLuc-ASARTDL secretion following hyperthermia and MDMA treatment (Fig. 5E-F).

**Figure 5:**
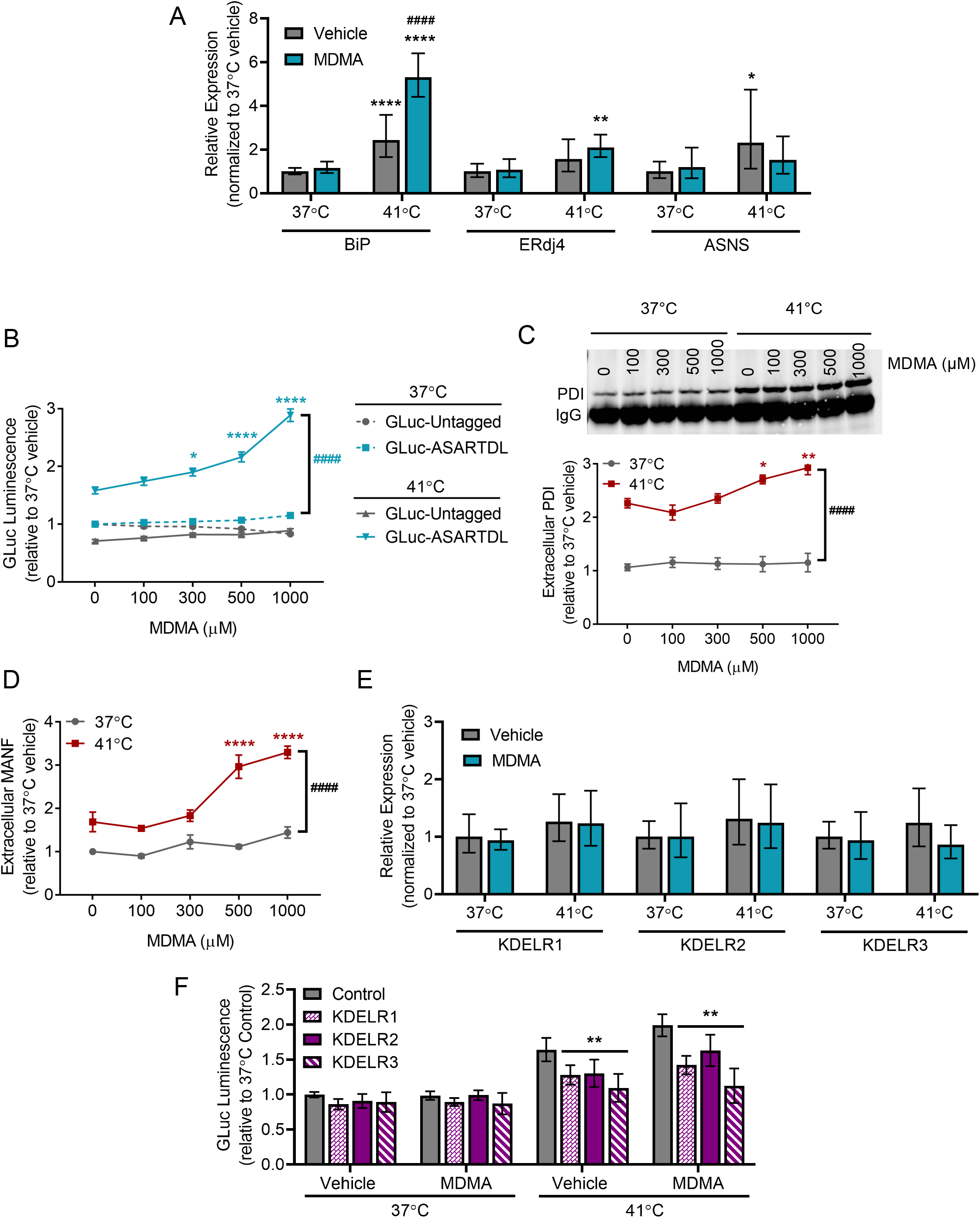
MDMA changes cellular responses to hyperthermia. (A) BiP, ERdj4, and ASNS mRNA levels analyzed with real-time RT-qPCR after treatment with vehicle or 1 mM MDMA and an 8 h exposure to 37°C or 41°C (2^−ddCq^ ± upper and lower limits, n=8, two-way ANOVA with Sidak’s multiple comparisons, *p<0.05, **p<0.01 and ****p<0.0001 37°C vs. 41°C, ^####^p<0.001 vehicle vs. caffeine). (B) GLuc activity in the media from SH-SY5Y cells stably expressing GLuc-ASARTDL or GLuc-Untagged after treatment with vehicle or MDMA and a 24 h incubation at 37°C or 41°C (mean ± SEM, n=26, two-way ANOVA with Tukey’s multiple comparisons, ^####^p<0.0001 37°C vs. 41°C, *p<0.05, ****p<0.0001 vehicle vs. caffeine). (C-D) Fold change in (C) immunoprecipitated PDI (representative blot shown) or (D) MANF in media from SH-SY5Y cells treated with vehicle or MDMA and incubated for 24 h at 37°C or 41°C (mean ± SEM, n=6, two-way ANOVA with Dunnett’s multiple comparisons, ^####^p<0.0001 37°C vs. 41°C, *p<0.05, **p<0.01, and ****p<0.0001 vehicle vs. caffeine). (E) KDELR1, KDELR2, and KDELR3 mRNA levels analyzed with real-time RT-qPCR after treatment with vehicle or 1 mM MDMA and an 8 h exposure to 37°C or 41°C (2^−ddCq^ ± upper and lower limits, n=8, two-way ANOVA with Sidak’s multiple comparisons). (F) GLuc activity in media following transduction of SH-SY5Y cells stably expressing GLuc-ASARTDL with lentiviral KDEL receptors and treatment with vehicle or 1 mM MDMA and a 24 h exposure to 37°C or 41°C (mean ± SEM, n=9, two-way ANOVA with Dunnett’s multiple comparisons, p<0.0001 37°C vs. 41°C, **p<0.01 control vs. LV-KDELR).

From a club drug perspective, the combination of caffeine, MDMA, and hyperthermia warrants further study. Clubs are often crowded venues with dancing both of which increase the ambient temperature of the room. MDMA is a common club drug and is often combined with caffeine through the form of drinks containing caffeine. MDMA is also one of the most adulterated drugs and caffeine is one of the substances used to cut MDMA. Treating SH-SY5Y with MDMA resulted in a leftward shift of a caffeine dose response with an increase in hyperthermia induced GLuc-ASARTDL secretion observed in cells treated with MDMA and 1 mM caffeine (Fig. 6A). At no dose of caffeine was extracellular GLuc-ASARTDL increased in the normal temperature control (37°C) regardless of whether the cells were treated with MDMA (Fig. 6A). Furthermore, no condition changed GLuc-Untagged secretion (Supp. Fig. 6A). As observed previously, increasing doses of caffeine diminished cellular ATP (Supp. Fig. 6B). Using 1 mM caffeine in combination with a dose response of MDMA revealed a significant potentiation of GLuc-ASARTDL hyperthermia-induced secretion with MDMA treatment (Fig. 6B). The combination of caffeine, MDMA, and hyperthermia also led to increased secretion of the endogenous ERS-containing protein MANF (Fig. 6C). No change in GLuc-Untagged secretion was observed and while higher doses of MDMA led to decreased cellular ATP the presence of caffeine did not significantly exacerbate ATP loss (Supp. Fig. 6C-D). No combination of temperature, MDMA, or caffeine increased KDEL receptor expression (Sup. Fig. 6D). While hyperthermia did alter UPR gene expression the combination of MDMA and caffeine did not cause BiP, ERdj4, or ASNS upregulation beyond that seen with either drug alone (Fig. 6E). Together these data indicate that caffeine and MDMA enhance ER resident protein secretion during hyperthermia and the combination of drugs further augments secretion. While MDMA increased BiP expression in hyperthermic conditions, other UPR markers are not affected by caffeine, MDMA, or a combination of the two drugs.

**Figure 6:**
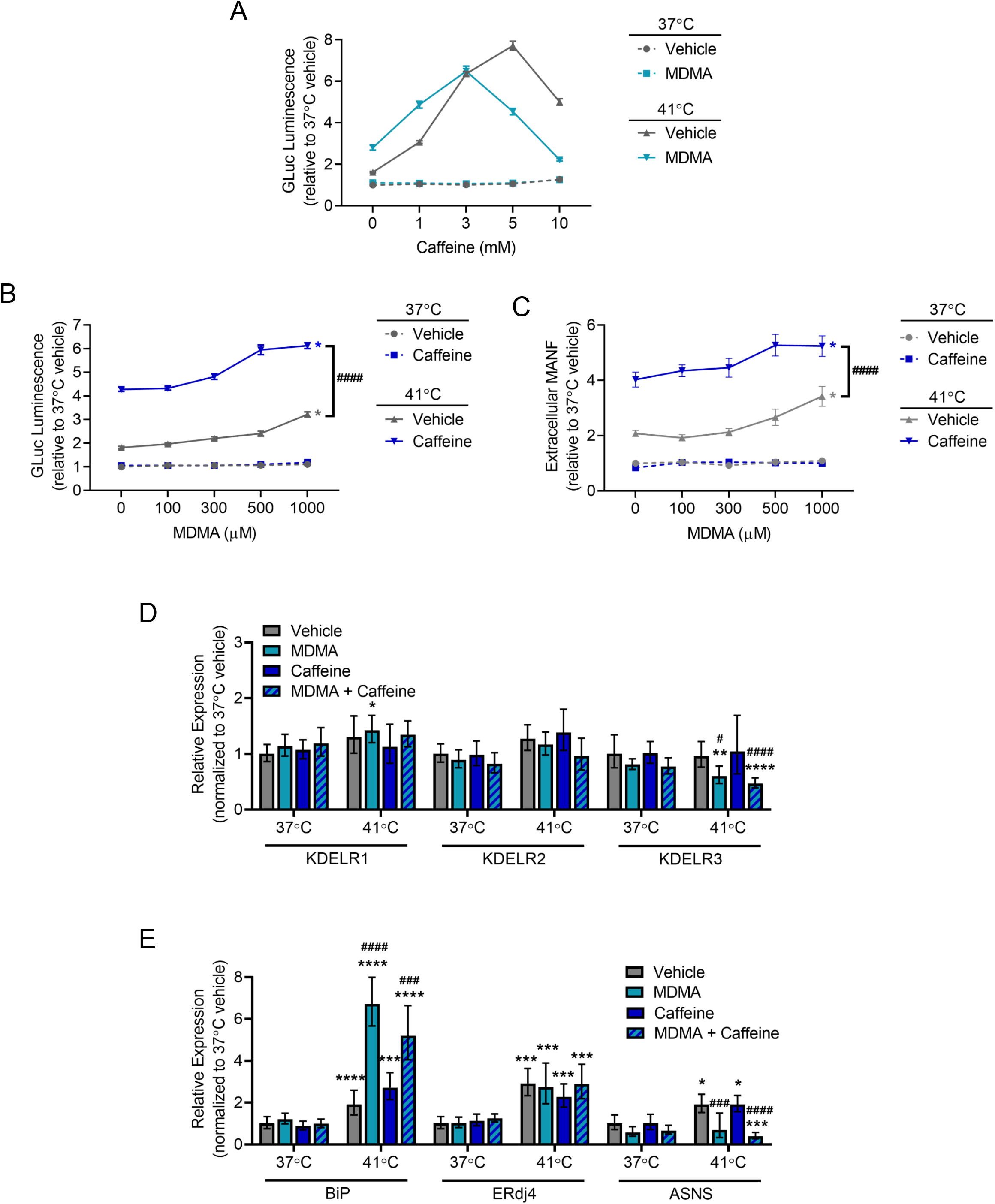
Caffeine and MDMA in combination affect hyperthermia-induced cellular responses. (A) GLuc activity in the media from SH-SY5Y cells stably expressing GLuc-ASARTDL after treatment with vehicle or 500 μM MDMA in combination with a dose response of caffeine (mean ± SEM, n=24). (B) GLuc activity in the media from SH-SY5Y cells stably expressing GLuc-ASARTDL after treatment with vehicle or 1 mM caffeine in combination with a dose response of MDMA and incubation at 37°C or 41°C (mean ± SEM, n=24, three-way ANOVA with Slice decomposition, *p<0.05 vehicle vs. MDMA at 41°C, ^####^p<0.0001 vehicle vs. caffeine at 41°C). (C) MANF in media from SH-SY5Y cells treated with vehicle or 1 mM caffeine in combination with a dose response of MDMA and incubation at 37°C or 41°C (mean ± SEM, n=6, three-way ANOVA with Slice decomposition, *p<0.05 vehicle vs. MDMA at 41°C, ^####^p<0.0001 vehicle vs. caffeine at 41°C. (D) KDELR1, KDELR2, and KDELR3 mRNA levels analyzed with real-time RT-qPCR after treatment with vehicle, 1 mM MDMA, 1 mM caffeine, or a combination of 1 mM MDMA and 1 mM caffeine and an 8 h exposure to 37°C or 41°C (2^−ddCq^ ± upper and lower limits, n=8, two-way ANOVA with Sidak’s multiple comparisons, *p<0.05, **p<0.01 and ****p<0.0001 37°C vs. 41°C, ^#^p<0.05 and ^####^p<0.001 vehicle vs. drug treatment). (E) BiP, ERdj4, and ASNS mRNA levels analyzed with real-time RT-qPCR after treatment with vehicle, 1 mM MDMA, 1 mM caffeine, or a combination of 1 mM MDMA and 1 mM caffeine and an 8 h exposure to 37°C or 41°C (2^−ddCq^ ± upper and lower limits, n=8, two-way ANOVA with Sidak’s multiple comparisons, *p<0.05, ***p<0.001 and ****p<0.0001 37°C vs. 41°C, ^####^p<0.001 and ^####^p<0.0001 vehicle vs. drug treatment).

## Discussion

The lumen of the endoplasmic reticulum is the site of many critical cellular functions such as protein folding and trafficking, lipid and carbohydrate metabolism, drug detoxification, and intracellular calcium storage. Resident proteins with ERS tails mediate these functions and are secreted when ER calcium is depleted in a process referred to as exodosis ^36^. The consequences of club drugs and hyperthermic conditions on ER proteostasis as it is related to the localization of ER resident proteins has not been previously reported. Here, we used both an exogenous reporter of ER exodosis, GLuc-ASARTDL, and endogenous ERS proteins (MANF, PDI, esterases) to show that hyperthermia triggered ER exodosis in both human and rodent neuronal cells. We also demonstrate that both caffeine and MDMA increase the magnitude of ER resident protein secretion under hyperthermic conditions.

Previous findings indicate that ER exodosis and GLuc-ASARTDL secretion is linked to depletions in ER calcium^35^. Calcium within the ER lumen is usually maintained at very high concentrations with estimations putting ER calcium concentrations at 1,000 to 10,000 times greater than in the cytosol ^54-58^. Hyperthermia causes an increase in intracellular free calcium that is thought to be at least partially dueto the release of calcium from internal stores like the ER^37^. Our results implicate a role for ER calcium in hyperthermia as drugs that stabilize ER calcium (dantrolene and 2-APB) by blocking ER calcium efflux channels reduced the ER protein secretion seen at elevated temperatures. We also found that further depleting ER calcium with thapsigargin during hyperthermic conditions potentiates ER exodosis. Changes in ER calcium stores may also underlie the enhanced ER exodosis seen in both caffeine and MDMA treated cells exposed to hyperthermia conditions. Caffeine is a documented RyR agonist that induces calcium release from intracellular ER calcium stores, while MDMA treatment is associated with increased cytosolic calcium levels resulting at least in part from depletions of intracellular ER calcium stores as blockers of ER calcium efflux (dantrolene and 2-APB) attenuate MDMA-induced cytosolic calcium increases ^59-61^. In addition to these direct links to ER calcium stores, caffeine and MDMA are also associated with indirect processes that may affect ER calcium. Caffeine is a phosphodiesterase (PDE) inhibitor that may affect ER calcium stores through cAMP signaling. PDE normally breaks down cAMP, but caffeine blocks this process leading to increased IP3-evoked ER calcium release^62^. MDMA stimulates serotonin release which works through a G-protein coupled receptor pathway in which phospholipase C (PLC) is activated and hydolyzes phosphatidylinositol biphosphate (PIP2) into IP3. IP3 can then stimulate ER calcium release through a second messenger system^63^. Our results showing increased ER resident protein secretion in hyperthermic conditions that was further enhanced by caffeine and MDMA supports the need for additional studies examining the specific interplay of ER calcium stores, hyperthermia, caffeine, and MDMA *in vivo*.

Although the exact mechanism by which hyperthermia triggers ER resident protein secretion remains unknown, changes in the unfolded protein response (UPR) and KDEL receptors may play a role in modulating this secretion. Prolonged exposure to hyperthermia (8 h and 24 h) was associated with upregulation of three UPR target genes. Further experimentation showed that the IRE1α pathway of the UPR was of particular importance as inhibiting IRE1α kinase activity potentiated extracellular GLuc-ASARTDL levels in hyperthermic conditions. When ERS-containing proteins leave the ER they interact with KDEL receptors in the Golgi and are transported back to the ER^25, 30^. We previously demonstrated that KDEL receptors are upregulated by ER stress following XBP1 activation ^36^. In our hyperthermia model, we saw an increase in ERdj4 expression (an XBP1 regulated gene) following hyperthermia and observed increased ER exodosis following treatment with KIRA6, a drug that prevents XBP1 activation by blocking IRE1α. Given the connection between XBP1 and KDEL receptors we hypothesized that KDELR2 and KDELR3 expression may be increased in hyperthermic conditions because those two isoforms of KDEL receptors are IRE1α/XBP1 regulated. However, we observed minimal KDEL receptor upregulation following hyperthermia. While we observed no change in endogenous KDEL receptor mRNA levels, manipulations of KDEL receptors did affect ER exodosis. Knockdown of KDEL receptors increased ER resident protein secretion in both normothermic and hyperthermic environments, while overexpression of KDEL receptors attenuated ER exodosis following hyperthermia. This makes KDEL receptor activation an attractive therapeutic target. Additional examinations of the UPR and KDEL receptor retrieval pathway is needed to determine precisely what role these processes play in hyperthermia and how they affect hyperthermia-induced ER exodosis. It is also worth noting that elevated temperatures change the cellular environment and are associated with decreased RNA, DNA, and protein synthesis, which may be one reason that no upregulation of KDEL receptors was observed ^64^.

The behavior of ER resident proteins at high temperatures also needs to be taken into consideration when experimental procedures require elevations in temperature. For example, a commonly used marker of exocytic membrane trafficking is temperature sensitive with a downshift in temperature inducing trafficking^65^. The high temperatures (around 40°C) that cells are held at prior to the induction of trafficking has the potential to lead to ER exodosis and ER stress, which could affect the subsequently observed trafficking process. Another instance in which hyperthermia triggered ER exodosis should be considered is in the use of temperature inducible promoters^66, 67^. Elevations in temperature trigger the expression of the gene of choice but could also prompt unintended changes in cellular proteostasis..

Taken together, our data support a model in which ER resident proteins, including some UPR proteins, are secreted from the cell in response to elevated temperatures. Our observations indicate that such a disruption of ER proteostasis may be a symptom of prolonged hyperthermia and could underlie hyperthermia associated drug toxicity. Currently, patients with elevated body temperature are given anti-pyretics or treated non-pharmacologically with body cooling procedures (e.g. ice, cooling blankets)^68^. However, these treatments do not directly address the cellular deficits that manifest following elevated temperatures. As ER exodosis represents a drug targetable cellular mechanism of dysfunction, our findings have implications in the treatment of hyperthermia associated impairments^69^. Furthermore, the convergence of elevated temperatures with caffeine or MDMA exposure to potentiate ER exodosis constitutes a therapeutic opportunity for decreasing drug-induced cellular toxicity associated with club drug usage. In case reports of MDMA overdose, treatment with dantrolene has been part of a successful treatment paradigm, and the ability of dantrolene to promote ER proteostasis may provide a cellular explanation for this success^70, 71^. Treatment options for substances of abuse remain limited and targeting ER exodosis in hyperthermia associated overdoses could assist in recovery. Alternatively, further augmenting the UPR with small molecular regulators of the UPR may aid in the recovery from acute overdose^72^.

## Acknowledgments

We thank Lowella Fortuno for preparing primary cortical neuron cultures, Doug Howard and the NIDA Genetic Engineering and Viral Vector Core for producing viral vectors, and David Epstein for his guidance regarding statistical tests. This work was supported by the Intramural Research Program at the National Institute on Drug Abuse.

## Methods

### Reagents

Dantrolene (Sigma), thapsigargin (Sigma), tunicamycin (Sigma), dithiothreitol (DTT; Sigma), KIRA6 (Cayman Chemicals), 2-Aminoethyl diphenylborinate (2-APB; Sigma), caffeine (Sigma), 3,4-methylenedioxymethamphetamine (DL-MDMA HCl, NIH/NIDA Pharmacy)

### SH-SY5Y Cell Culture

The creation of SH-SY5Y cells stably expressing either GLuc-ASARTDL or GLuc-Untagged was described previously^35^. Cells were maintained in Dulbecco’s Modified Eagle Medium (DMEM) + GlutaMAX, 4.5 g/L D-glucose, 110 mg/L sodium pyruvate (Thermo Fisher Scientific) supplemented with 10% bovine growth serum (GE Life Sciences), 10 units/mL penicillin, and 10 μg/mL streptomycin (Thermo Fisher Scientific). Cells were grown at 37°C with 5.5% CO_2_ in a humidified incubator. Cells were plated in 96-well plates with 5 × 10^4^ cells/well or 24-well plates with 2.5 × 10^5^ cells/well. After 24 h, cells were either transferred to an incubator maintained at 41°C or remained at 37°C. Incubator temperatures were monitored with a K-type thermocouple probe (Omega).

### Primary Cortical Neuron Cell Culture

Rat primary cortical neurons were prepared from Sprague-Dawley rats on embryonal day 15 (E15) as previously described and in accordance with National Institutes of Health Animal Care and Usage Committee guidelines ^73^. Embryos of both sexes were used, and cells were plated on polyethyleneimine-coated 96-well plates with 6.0 × 10^4^ cells/well. Cells were maintained in neurobasal media (Thermo Fisher Scientific) supplemented 2% B-27 (Thermo Fisher Scientific) and 0.5 mM L-glutamine (Sigma) at 37°C with 5.5% CO_2_ in a humidified incubator. A 50% media exchange was done on days 4, 6, 8, 11, and 13. Viral transductions were done on day 6 with 5 μL of AAV1-CaMKII-GLuc-ASARTDL (Addgene 149503) or AAV1-CaMKII-GLuc-Untagged (Addgene 149502) at 5.48 × 10^9^ vg/mL. On day 13, cells were either transferred to an incubator maintained at 41°C or remained at 37°C. Incubator temperatures were monitored with a K-type thermocouple probe (Omega).

### Gaussia Luciferase Assay

*Gaussia* luciferase (GLuc) assays were performed as described previously^35^. Briefly, 5 μL of cell culture media was transferred to an opaque walled plate and luciferase levels were determined using a Biotek Synergy 2 plate reader with an injector setup. Following an injection of 100 μL of 10 μM coelenterazine (Regis Technologies) luminescence at 25°C was measured with a sensitivity of 100, a 0.5 sec integration time, and a 5 sec delay.

### ATP Assay

The Promega CellTiter-Glo Luminescent Cell Viability Assay (ATP assay) was used as per the manufacturer’s protocol. Briefly, an equal volume of ATP substrate was added to the cell culture plate and incubated with agitation for 10 min at room temperature. After a further 2 min incubation without agitation, 100 μL of solution was transferred to an opaque walled plate and luminescence was read using a Biotek Synergy 2 plate reader.

### PDI Immunoprecipitation

100 μL of protein A beads (SureBeads, Bio-Rad) were washed with PBS + 0.1% Tween-20 (PBS-T) then incubated with PDI antibody (Abcam, ab2792) diluted 1:100 in200 μL PBS-T for 10 min. Beads were again washed with PBS-T then incubated with 400 μL cell culture media for 1 h. Following a final washing with PBS-T, samples were eluted with 40 μL of 1x LDS (Thermo Fisher Scientific). After heating samples at 70°C for 10 min, equal volumes of sample were loaded into a 4-12% Bis-Tris gel (Thermo Fisher Scientific) and run using 1x MOPS buffer (Thermo Fisher Scientific). An iBlot2 (Thermo Fisher Scientific) was used to transfer proteins to a 0.2 μm PVDF membrane (Thermo Fisher Scientific) using Program P0. Blots were blocked at room temperature for 1 h using Rockland blocking buffer (VWR). Primary PDI antibody (Abcam, ab2792) diluted 1:500 was applied overnight at 4°C and secondary goat-anti mouse IR680 antibody (LICOR) diluted 1:4000 was applied for 1 h at room temperature. Blots were scanned using an Odyssey scanner (LICOR).

### MANF Homogeneous Time Resolved Fluorescence (HTRF) Assay

The amount of MANF present in cell culture media samples was quantified using a MANF HTRF assay (Cisbio) as per the manufacturer’s instructions. Briefly, media samples diluted 1:2 in dilution buffer were incubated with anti-Human MANF-d2 antibody and anti-Human-Eu^3+^ cryptate antibody for 24 h.

Fluorescence was measured at 665 nm and 620 nm and sample readouts were compared to a MANF standard curve.

### Esterase Assay

Esterase activity was measured as previously described^34^. Immediately before hyperthermia treatments began cells underwent a full media exchange into esterase assay media (150 mM NaCl, 5 mM KCl, 1 mM MgCl_2_, 20 mM HEPES, 1 mM CaCl_2_, and 1.9 g/L glucose). An interaction between esterases and a fluorescein di-(1-methylcyclopropanecarboxymethyl ether) produces fluorescence that was quantified by a BioTek Synergy H1 plate reader (Excitation 465 nm/Emission 528 nm). An equal volume of cell culture media and esterase substrate (100 μM in esterase assay media at pH 5) was transferred to a black walled clear bottomed plate (Perkin Elmer) and fluorescence was measured after 1 h.

### siRNA Transfection

SH-SY5Y cells (2.5 × 10^4^ cells/well, 96-well plate) stably expressing GLuc-ASARTDL or GLuc-Untagged were reverse transfected with 10 nM siRNA using Lipofectamine RNAiMax (Thermo Fisher Scientific). A full media exchange was performed 48 h after transfection and after a further 24 h incubation cells were placed in hyperthermic or normothermic conditions for 24 h. The siRNA used were from the Thermo Fisher Scientific Silencer Select Library: KDELR1 (assay s548), KDELR2 (assay s21689), KDELR3 (assay s21690), control (Cat. #4390843).

### Lentiviral Transduction

Lentiviral vectors expressing Myc and FLAG tagged KDEL receptors 1, 2 and 3 were previously described^33^. Viruses were titered using the Lenti-X p24 rapid titer kit (Takara). SH-SY5Y were transduced in 96-well plates at multiplicity of infection (MOI) of 2, incubated for 48 h, and then exposed to hyperthermic conditions as described above.

### Real time qPCR

RNA was isolated from cells using a NucleoSpin RNA Plus kit (Macherey-Nagel) according to the manufacturer’s protocol. Using iScript (Bio-Rad) 500 ng of RNA was transcribed into cDNA in a 20 μL reaction. The cDNA was diluted 1:40 with DNase-free water. 5 μL of cDNA was assayed in duplicate in a 20 μL reaction composed of TaqMan Universal PCR Master Mix (Thermo Fisher Scientifc), 450 nM primers, and 100 nM probe. Real time qPCR was performed using a C1000 Thermal Cycler CFX96 Real-Time System (Bio-Rad) with a pre-incubation (50°C for 10 s followed by photobleach, repeat 20x, 95°C for 5 min), and amplification with 50 repeats (94°C for 20 s, 60°C for 1 min). All C_q_ values were normalized to the C_q_ for glyceraldehyde 3-phosphate dehydrogenase (GAPDH). Two other reference genes, Ubiquitin-conjugating enzyme 2i (Ube2i) and RNA polymerase II (PRNAII), were evaluated for use, but did not remain constant over treatment paradigms. Results are presented as the 2^-ddCq^ value ± upper and lower limits (limits calculated based on the standard deviation of delta Cq values). The primer and probe sequences used were: human <u>ASNS,</u>ggattggctgccttttatcagg (forward), ggcttctttcagctgcttcaac (reverse), tggactccagcttggttgctgcc (FAM-labeled probe); human BiP, gttgtggccactaatggagatac (forward), ggagtttctgcacagctctattg (reverse), acgctggtcaaagtcttctccaccca (FAM-labeled probe); human ERdj4, gccatgaagtaccaccctg (forward), ccactagtaaaagcactgtgtc (reverse), ctgcaatctctctgaattttgcttcagc (FAM-labeled probe); human PRNAII, gcaccacgtccaatgacattg (forward), ggagccatcaaaggagatgac (reverse), acggcttcaatgcccagcaccg (HEX-labeled probe); human Ube2i, gtgtgcctgtccatcttagag (forward), gctgggtcttggatatttggttc (reverse), caaggactggaggccagccatcac (HEX-labeled probe); human KDELR1, tgctccttcaccacggtctg (forward), ggtgaagtcatgattgaccaggaacg (reverse), agttcctggtcgttcccacagccattctg (FAM-labeled probe); human KDELR2, ctatgccacagtgtacctgatc (forward), agagaaatcgtgattaactaaaaatgag (reverse), ctacgatggaaatcatgataccttccgag (FAM-labeled probe); human KDELR3, gaggtccaagtgctgcaagg (forward), cactgtaacataggcacagaggag (reverse), caccaggtacctggacctgttcaccaa (FAM-labeled probe) (Integrated DNA Technologies); human GAPDH, CATCATCCCTGCCTCTACTG (forward), CTTGCCCACAGCCTTGGCAGC (reverse), CCAGTGAGCTTCCCGTTCA (HEX-labeled probe).

## Figure Legends

**Supplemental Figure 1:**
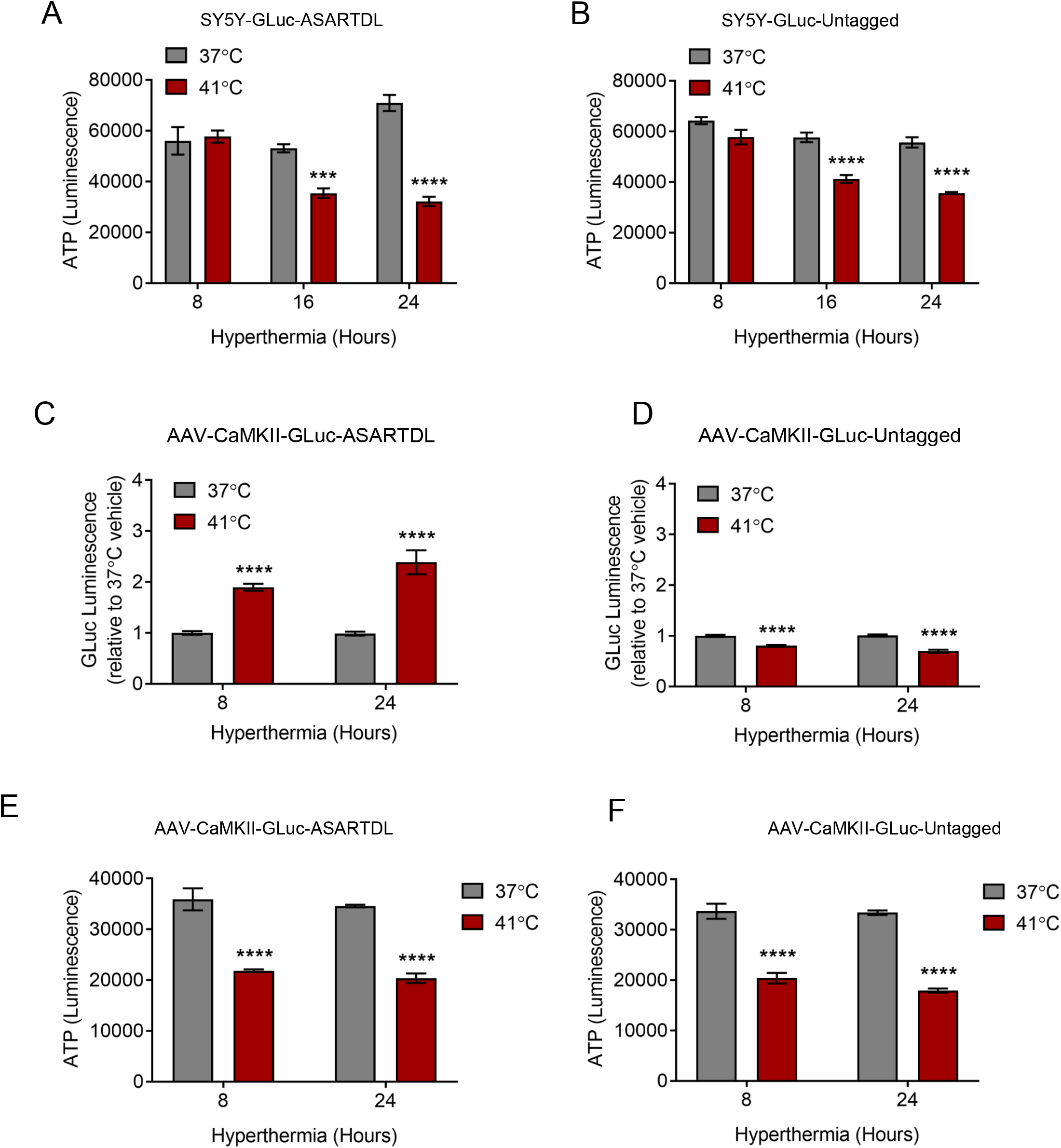
Hyperthermia increases GLuc-ASARTDL secretion and decreases cell metabolic activity. (A-B) ATP assay of SH-SY5Y cells stably expressing (A) GLuc-ASARTDL or (B) GLuc-Untagged after an 8 h, 16 h, or 24 h incubation at 37°C or 41°C (mean ± SEM, n=6, two-way ANOVA with Sidak’s multiple comparisons, ***p<0.001 and ****p<0.0001 37°C vs. 41°C). (C-D) GLuc activity in the media from PCNs transduced with (C) GLuc-ASARTDL or (D) GLuc-Untagged after an 8 h or 24 h incubation at 37°C or 41°C (mean ± SEM, n=27, two-way ANOVA with Sidak’s multiple comparisons, ****p<0.0001 37°C vs. 41°C). (E-F) ATP assay of PCNs transduced with (E) GLuc-ASARTDL or (F) GLuc-Untagged after an 8 h or 24 h incubation at 37°C or 41°C (mean ± SEM, n=6, two-way ANOVA with Sidak’s multiple comparisons, ****p<0.0001 37°C vs. 41°C).

**Supplemental Figure 2:**
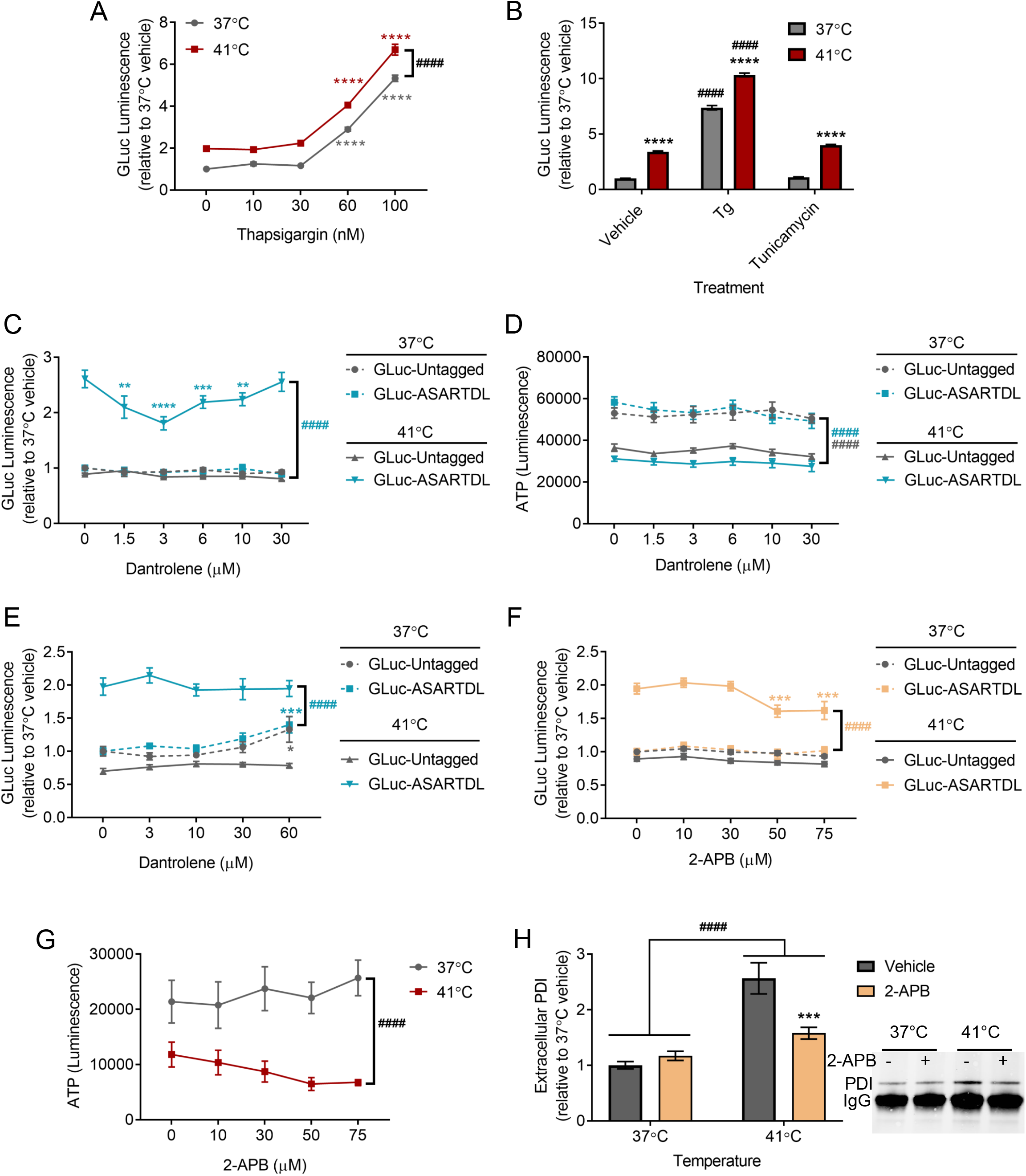
Modulation of ER calcium affects hyperthermia-induced ER exodosis. (A) GLuc activity in the media from SH-SY5Y cells stably expressing GLuc-ASARTDL following treatment with vehicle or thapsigargin and a 24 h incubation at 37°C or 41°C (mean ± SEM, n=16, two-way ANOVA with Dunnett’s multiple comparisons, ^####^p<0.0001 37°C vs. 41°C, ****p<0.0001 vehicle vs. thapsigargin). (B) GLuc activity in the media from SH-SY5Y cells stably expressing GLuc-ASARTDL following treatment with vehicle, 200 nM thapsigargin, or 3 μg/mL tunicamycin treatment and a 24 h incubation at 37°C or 41°C (mean ± SEM, n=24, two-way ANOVA with Tukey’s multiple comparisons, ****p<0.0001 37°C vs. 41°C, ^####^p<0.0001 vehicle vs. drug). (C) GLuc activity in the media from SH-SY5Y cells stably expressing either GLuc-ASARTDL or GLuc-Untagged after a 16 h pre-treatment with dantrolene followed by a 24 h incubation at 37°C or 41°C (mean ± SEM, n≥9, two-way ANOVA with Dunnett’s multiple comparisons, ^####^p<0.0001 37°C vs. 41°C, **p<0.01, ***p<0.001, and ****p<0.0001 vehicle vs. dantrolene) (D) ATP assay of SH-SY5Y cells stably expressing GLuc-ASARTDL or GLuc-Untagged after a 16 h pre-treatment with dantrolene followed by a 24 h incubation at 37°C or 41°C (mean ± SEM, n=9, two-way ANOVA with Dunnett’s multiple comparisons, ^####^p<0.0001 37°C vs. 41°C). (E) GLuc activity in the media from PCNs transduced with either GLuc-ASARTDL or GLuc-Untagged after a 30 min pre-treatment with dantrolene followed by a 24 h incubation at 37°C or 41°C (mean ± SEM, n=6, two-way ANOVA with Dunnett’s multiple comparisons, ^####^p<0.0001 37°C vs. 41°C, ***p<0.001 vehicle vs. dantrolene). (F) GLuc activity in the media from PCNs transduced with either GLuc-ASARTDL or GLuc-Untagged after a 30 min pre-treatment with 2-APB followed by a 24 h incubation at 37°C or 41°C (mean ± SEM, n=18, two-way ANOVA with Dunnett’s multiple comparisons, ^####^p<0.0001 37°C vs. 41°C, ***p<0.001 vehicle vs. 2-APB). (G) ATP assay of PCNs after a 30 min pre-treatment with 2-APB followed by a 24 h incubation at 37°C or 41°C (mean ± SEM, n=9, two-way ANOVA with Dunnett’s multiple comparisons, ^####^p<0.0001 37°C vs. 41°C). (H) Fold change in immunoprecipitated PDI (representative blot shown) in media from PCNs pre-treated with vehicle or 50 μM 2-APB for 30 min then incubated for 24 h at 37°C or 41°C (mean ± SEM, n=6, two-way ANOVA with Sidak’s multiple comparisons, ^####^p<0.0001 37°C vs. 41°C, ***p<0.001 vehicle vs. 2-APB).

**Supplemental Figure 3:**
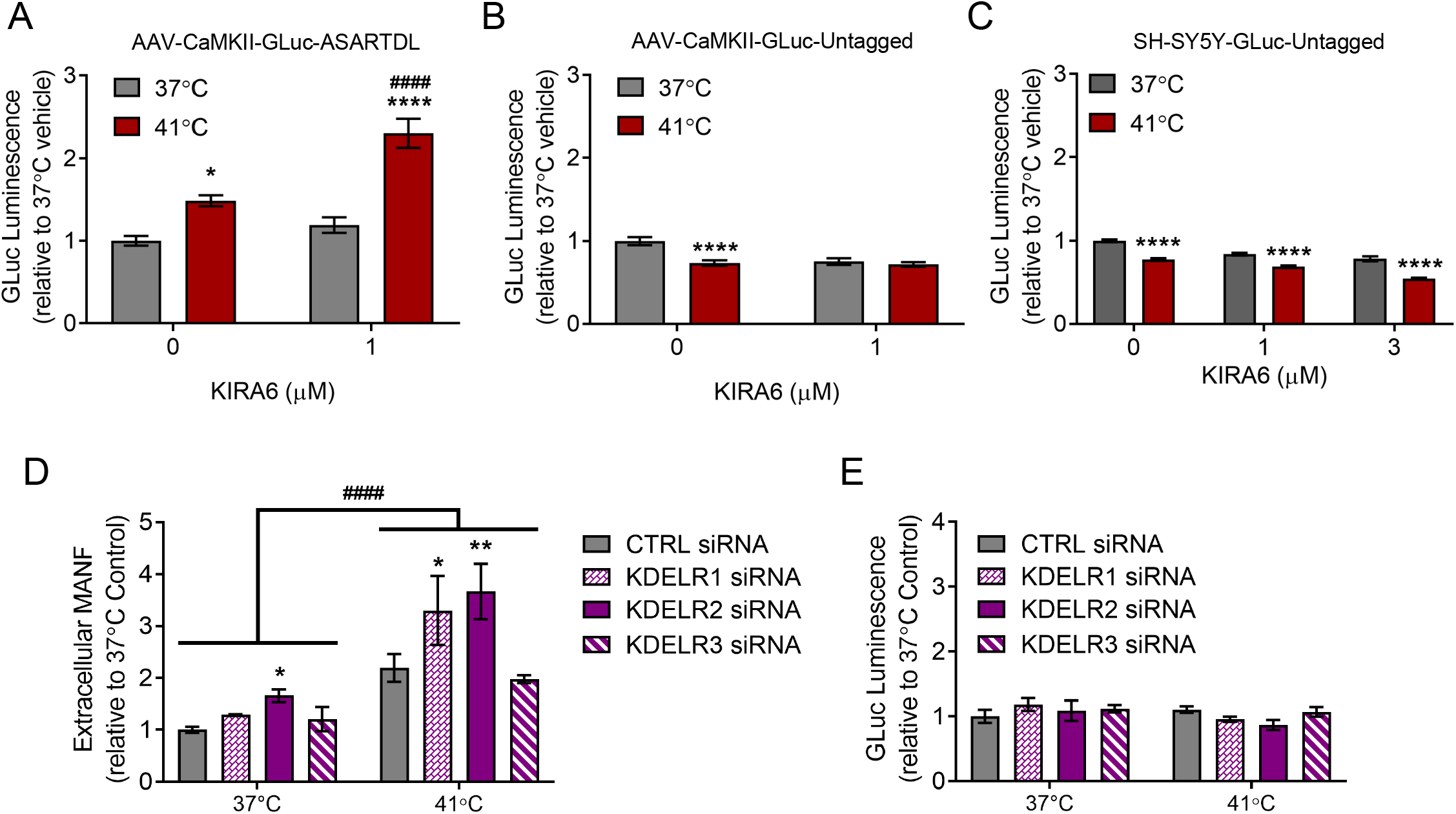
Hyperthermia associated changes related to the UPR and KDEL receptors. (A-B) GLuc activity in the media from PCNs transduced with (A) GLuc-ASARTDL or (B) GLuc-Untagged after a 1 h pre-treatment with vehicle or 1 μM KIRA6 followed by a 24 h incubation at 37°C or 41°C (mean ± SEM, n=15, two-way ANOVA with Tukey’s multiple comparisons, *p<0.05 and ****p<0.0001 37°C versus 41°C, ^####^p<0.0001 vehicle vs. KIRA6). (C) GLuc activity in the media from SH-SY5Y cells stably expressing GLuc-Untagged after a 1 h pre-treatment with vehicle or KIRA6 (1 μM or 3 μM) followed by a 24 h incubation at 37°C or 41°C (mean ± SEM, n=48, two-way ANOVA with Tukey’s multiple comparisons, ****p<0.0001 37°C vs. 41°C). (D) MANF in media following transfection of SH-SY5Y with 10 nM KDEL receptor siRNA and a 24 h incubation at 37°C or 41°C (mean ± SEM, n=6, two-way ANOVA with Dunnett’s multiple comparisons, ^####^p<0.0001 37°C vs. 41°C, *p<0.05 and **p<0.01 control vs. KDELR siRNA). (E) GLuc activity in media following transfection of SH-SY5Y cells stably expressing GLuc-Untagged with 10 nM KDEL receptor siRNA and a 24 h incubation at 37°C or 41°C (mean ± SEM, n=12, two-way ANOVA with Dunnett’s multiple comparisons).

**Supplemental Figure 4:**
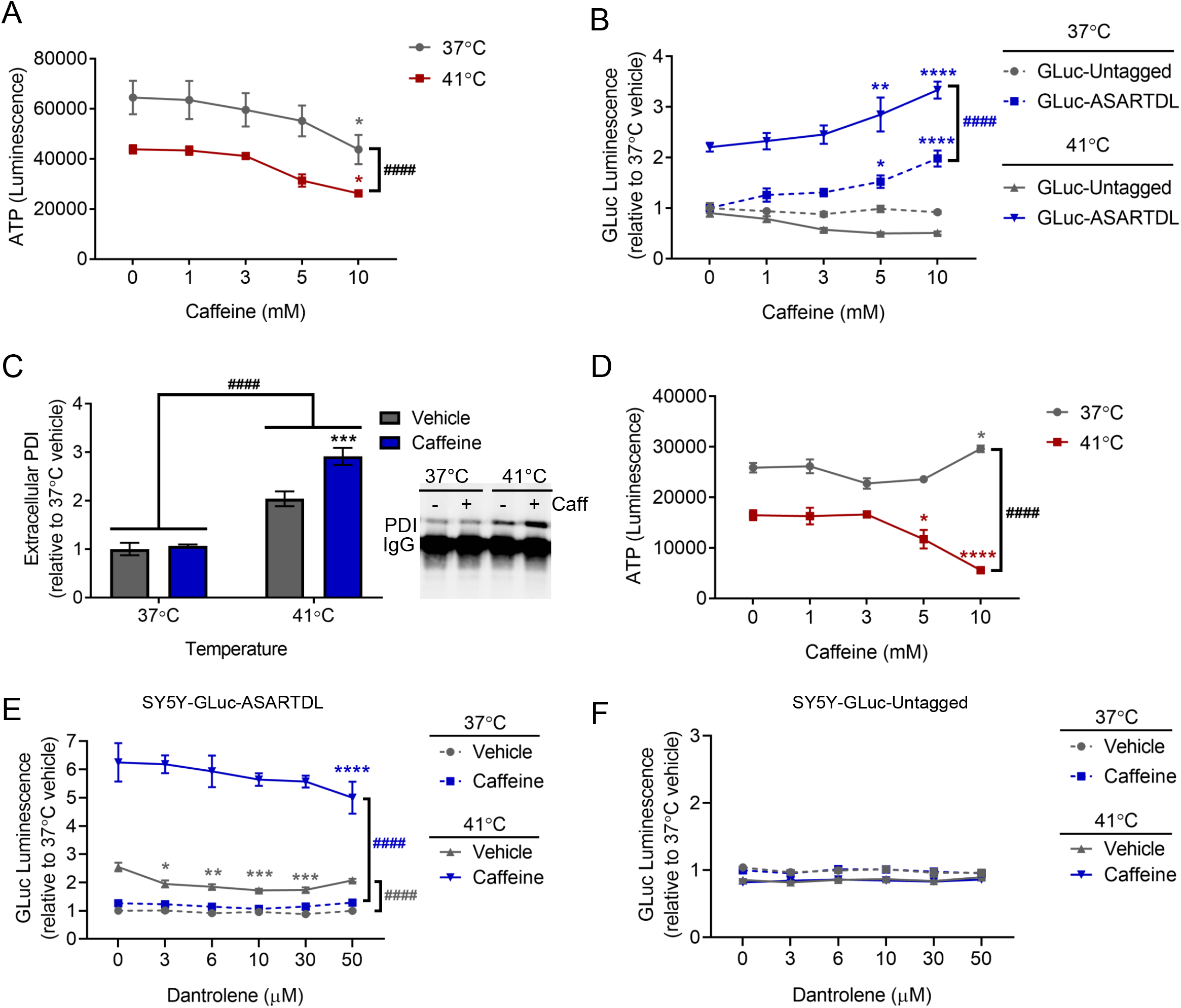
Caffeine associated changes in hyperthermia-induced ER. (A) ATP assay of SH-SY5Y cells stably expressing GLuc-ASARTDL after treatment with vehicle or caffeine and a 24 h incubation at 37°C or 41°C (mean ± SEM, n=6, two-way ANOVA with Dunnett’s multiple comparisons, ^####^p<0.0001 37°C versus 41°C, *p<0.05 vehicle vs. caffeine). (B) GLuc activity in the media from PCNs transduced with GLuc-ASARTDL or GLuc-Untagged after treatment with vehicle or caffeine and a 24 h incubation at 37°C or 41°C (mean ± SEM, n=12, two-way ANOVA with Dunnett’s multiple comparisons, ^####^p<0.0001 37°C vs. 41°C, *p<0.05, **p<0.01, ****p<0.0001 vehicle vs. caffeine). (C) Fold change in immunoprecipitated PDI (representative blot shown) in media from PCNs treated with vehicle or 5 mM caffeine and incubated for 24 h at 37°C or 41°C (mean ± SEM, n=6, two-way ANOVA with Sidak’s multiple comparisons, ^####^p<0.0001 37°C vs. 41°C, ***p<0.001 vehicle vs. caffeine). (D) ATP assay of PCNs transduced with GLuc-ASARTDL after treatment with vehicle or caffeine and a 24 h incubation at 37°C or 41°C (mean ± SEM, n=6, two-way ANOVA with Dunnett’s multiple comparisons, ^####^p<0.0001 37°C vs. 41°C, *p<0.05 and ****p<0.0001 vehicle vs. caffeine). (E) GLuc activity in the media from SH-SY5Y cells stably expressing GLuc-ASARTDL after a 16 h pre-treatment with dantrolene followed by treatment with 5 mM caffeine and a 24 h incubation at 37°C or 41°C (mean ± SEM, n=16, two-way ANOVA with Dunnett’s multiple comparisons, ^####^p<0.0001 37°C vs. 41°C, *p<0.05, **p<0.01, ***p<0.001, and ****p<0.0001 vehicle vs. dantrolene). (F) GLuc activity in the media from SH-SY5Y stably expressing GLuc-Untagged after a 16 h pre-treatment with dantrolene followed by treatment with 5mM caffeine and a 24 h incubation at 37°C or 41°C (mean ± SEM, n=16, two-way ANOVA with Dunnett’s multiple comparisons).

**Supplemental Figure 5:**
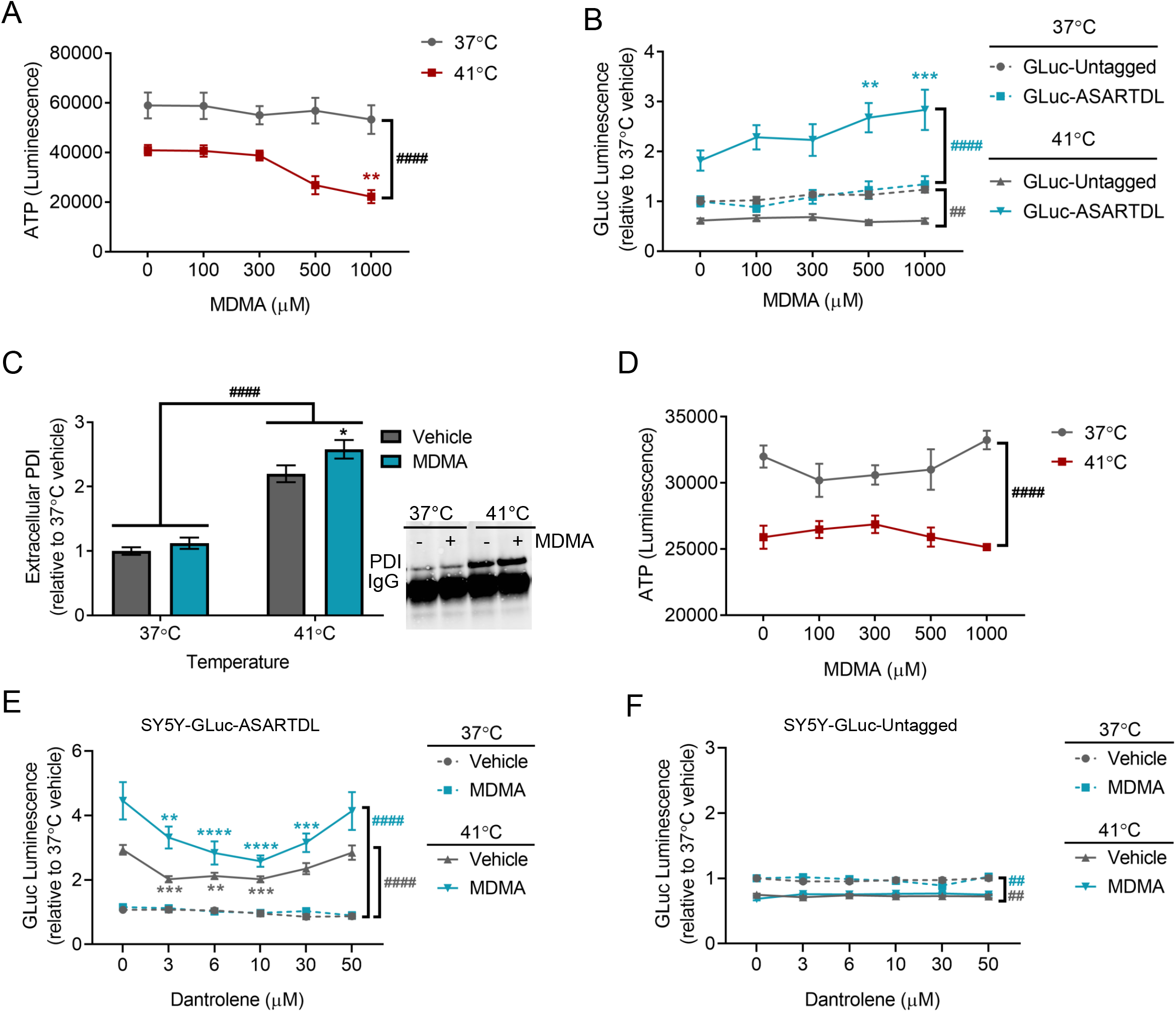
MDMA associated changes in hyperthermia-induced ER exodosis. (A) ATP assay of SH-SY5Y cells stably expressing GLuc-ASARTDL after treatment with vehicle or MDMA and a 24 h incubation at 37°C or 41°C (mean ± SEM, n=6, two-way ANOVA with Dunnett’s multiple comparisons, ^####^p<0.0001 37°C versus 41°C, **p<0.01 vehicle vs. MDMA). (B) GLuc activity in the media from PCNs transduced with GLuc-ASARTDL or GLuc-Untagged after treatment with vehicle or MDMA and a 24 h incubation at 37°C or 41°C (mean ± SEM, n=12, two-way ANOVA with Dunnett’s multiple comparisons, ^##^p<0.01 and ^####^p<0.0001 37°C vs. 41°C, **p<0.01 and ***p<0.001 vehicle vs. MDMA). (C) Fold change in immunoprecipitated PDI (representative blot shown) in media from PCNs treated with vehicle or 500 μM MDMA and incubated for 24 h at 37°C or 41°C (mean ± SEM, n=6, two-way ANOVA with Sidak’s multiple comparisons, ^####^p<0.0001 37°C vs. 41°C, *p<0.05 vehicle vs. MDMA). (D) ATP assay of PCNs transduced with GLuc-ASARTDL after treatment with vehicle or MDMA and a 24 h incubation at 37°C or 41°C (mean ± SEM, n=6, two-way ANOVA with Dunnett’s multiple comparisons, ^####^p<0.0001 37°C vs. 41°C). (E) GLuc activity in the media from SH-SY5Y cells stably expressing GLuc-ASARTDL after a 16 h pre-treatment with dantrolene followed by treatment with 1 mM MDMA and a 24 h incubation at 37°C or 41°C (mean ± SEM, n=16, two-way ANOVA with Dunnett’s multiple comparisons, ^####^p<0.0001 37°C vs. 41°C, **p<0.01, ***p<0.001, ****p<0.0001 vehicle vs. dantrolene). (F) GLuc activity in the media from SH-SY5Y stably expressing GLuc-Untagged after a 16 h pre-treatment with dantrolene followed by treatment with 1 mM MDMA and a 24 h incubation at 37°C or 41°C (mean ± SEM, n=16, two-way ANOVA with Dunnett’s multiple comparisons, ^##^p<0.01 37°C vs. 41°C).

**Supplemental Figure 6:**
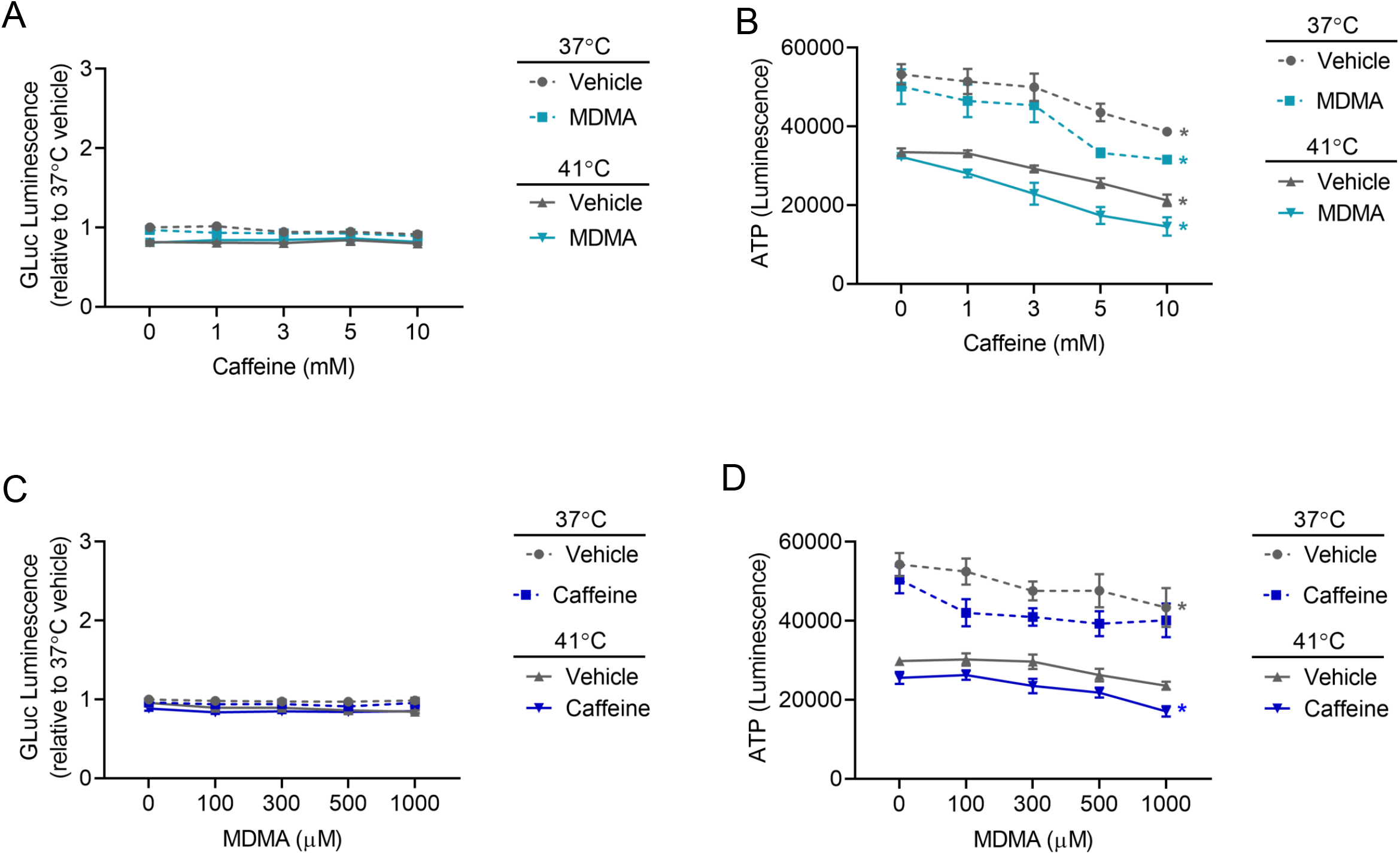
Caffeine and MDMA in combination affect cellular ATP, but not GLuc-Untagged secretion in hyperthermic conditions. (A) GLuc activity in the media from SH-SY5Y cells stably expressing GLuc-Untagged after treatment with vehicle or 500 μM MDMA in combination with a dose response of caffeine and a 24 h incubation at 37°C or 41°C (mean ± SEM, n=9, three-way ANOVA with Slice decomposition). (B) ATP assay of SH-SY5Y-GLuc-ASARTDL cells treated with vehicle or 500 μM MDMA in combination with a dose response of caffeine and a 24 h incubation at 37°C or 41°C (mean ± SEM, n=9, three-way ANOVA with Slice decomposition, p<0.001 37°C vs. 41°C, *p<0.05 vehicle vs. caffeine). (C) GLuc activity in the media from SH-SY5Y cells stably expressing GLuc-Untagged after treatment with vehicle or 1 mM caffeine in combination with a dose response of MDMA and a 24 h incubation at 37°C or 41°C (mean ± SEM, n=9, three-way ANOVA with Slice decomposition). (D) ATP assay of SH-SY5Y-GLuc-ASARTDL cells treated with vehicle or 1 mM caffeine in combination with a dose response of MDMA and a 24 h incubation at 37°C or 41°C (mean ± SEM, n=9, three-way ANOVA with Slice decomposition, p<0.001 37°C vs. 41°C, *p<0.05 vehicle vs. MDMA).

